# Mixed model association for biobank-scale data sets

**DOI:** 10.1101/194944

**Authors:** Po-Ru Loh, Gleb Kichaev, Steven Gazal, Armin P Schoech, Alkes L Price

**Author notes:** Correspondence should be addressed to P.-R.L. or A.L.P.

## Abstract

Biobank-based genome-wide association studies are enabling exciting insights in complex trait genetics, but much uncertainty remains over best practices for optimizing statistical power and computational efficiency in GWAS while controlling confounders. Here, we introduce a much faster version of our BOLT-LMM Bayesian mixed model association method— capable of running analyses of the full UK Biobank cohort in a few days on a single compute node—and show that it produces highly powered, robust test statistics when run on all 459K European samples (retaining related individuals). When used to conduct a GWAS for height in UK Biobank, BOLT-LMM achieved power equivalent to linear regression on 650K samples—a 93% increase in effective sample size versus the common practice of analyzing unrelated British samples using linear regression (UK Biobank documentation; Bycroft et al. bioRxiv). Across a broader set of 23 highly heritable traits, the total number of independent GWAS loci detected increased from 5,839 to 10,759, an 84% increase. We recommend the use of BOLT-LMM (retaining related individuals) for biobank-scale analyses, and we have publicly released BOLT-LMM summary association statistics for the 23 traits analyzed as a resource for all researchers.

## To the Editor

Despite recent work highlighting the advantages of linear mixed model (LMM) methods for genome-wide association studies in data sets containing relatedness or population structure [1–3], much uncertainty remains about best practices for optimizing GWAS power while controlling confounders. Several recent studies of the interim UK Biobank data set [4] (~ 150,000 samples) removed >20% of samples by applying filters for relatedness or genetic ancestry, and/or used linear regression in preference to mixed model association. These issues are exacerbated in the full UK Biobank data set (~500,000 samples), in which suggested sample exclusions decrease sample size by nearly 30% [5]. Here, we release a much faster version of our BOLT-LMM Bayesian mixed model association method [3] and show that it can be applied with minimal sample exclusions and achieves greatly superior power compared to common practices for analyzing UK Biobank data.

In analyses of 23 highly heritable UK Biobank phenotypes (Supplementary Table 1), we observed that BOLT-LMM (applied to all 459,327 European samples and ~20 million imputed variants passing QC) consistently achieved far greater association power than linear regression with principal component (PC) covariates (on 337,539 unrelated British samples, following ref. [5]), attaining an 84% increase in GWAS locus discovery (10,759 total independent loci versus 5,839; Fig. 1a and Supplementary Table 2). These gains in power were driven only partially by the increased number of samples analyzed; we observed that BOLT-LMM achieved effective sample sizes as high as ~700,000 by conditioning on polygenic predictions from genome-wide SNPs, which effectively reduces noise in an association test [2, 3, 6] (Fig. 1b, Supplementary Fig. 1, and Supplementary Table 3). (We estimated effective sample size by taking ratios of chi-square statistics computed by BOLT-LMM vs. linear regression at genome-wide significant SNPs; Supplementary Note.) The large sample size of the UK Biobank—which enables BOLT-LMM to predict and condition away up to 43% of phenotypic variance (approaching h_g_^2^ for several traits; Fig. 1b)—is now revealing the full power of this approach. We also confirmed that BOLT-LMM achieved substantial gains in power when run on the unrelated British sample set (Supplementary Tables 2 and 3).

**Figure 1.**
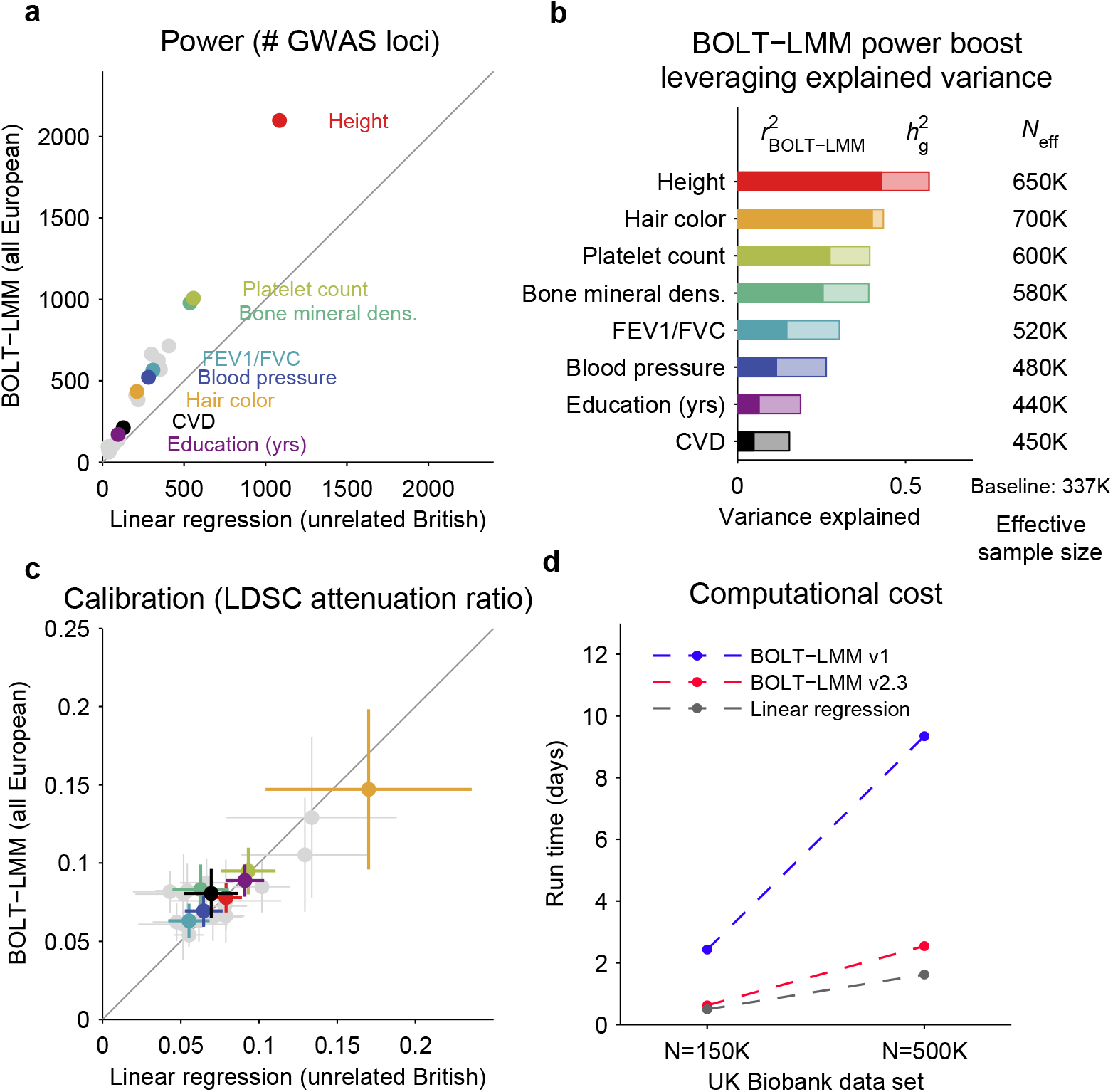
Power, calibration, and speed of BOLT-LMM v2.3 in UK Biobank analyses. **(a)** Numbers of independent genome-wide significant associations (*p*<5× 10^−9^) identified by BOLT-LMM analyses of all European-ancestry individuals (N=459,327) versus linear regression analyses of unrelated British individuals (*N*=337,539, following common practice [5]). Results for 23 phenotypes are plotted, with 8 representative phenotypes highlighted. **(b)** Variance explained by genome-wide SNPs on which BOLT-LMM implicitly conditions to increase power. Conditioning on BOLT-LMM’s polygenic predictions—which attain accuracy (*r*^2^_BOLT-LMM_) approaching SNP-heritability (*h*_*g*_^2^) for some traits—achieves effective sample sizes as high as ~700K. (We measured effective sample size by comparing *χ*^2^ statistics at associated SNPs; Supplementary Note.) **(c)** Test statistic calibration of BOLT-LMM on all European individuals versus linear regression on unrelated British individuals (using 20 principal component covariates). Attenuation ratios from LD score regression [7,8] match closely between the two methods, indicating that BOLT-LMM properly controls false positives (Supplementary Fig. 2). Error bars, jackknife s.e. **(d)** Computational cost of association analysis using BOLT-LMM v2.3, the previous version of BOLT-LMM [3], and linear regression (implemented efficiently within the BOLT-LMM software) on the UK Biobank N=150K and N=500K data releases. Analyses were run on 8 threads on a 2.10 GHz Intel Xeon E5-2683 v4 processor. Additional details and numerical data are provided in the Supplementary Note, Supplementary Fig. 1, and Supplementary Tables 1–7.

To verify that BOLT-LMM analyses of all European samples were robust to potential confounding due to relatedness or population structure, we performed LD score regression (LDSC) analyses [7] of association statistics computed using both BOLT-LMM (on all European samples—allowing related individuals as well as population structure) and linear regression (on unrelated British samples); we ran LDSC using the baselineLD model [8] using LD estimated in 1000 Genomes EUR samples. We observed that while the value of the LD score regression intercept (previously proposed as an indicator of confounding [7]) was generally difficult to interpret due to attenuation bias [3], which causes the intercept to rise above 1 with increased sample size and heritability (Supplementary Fig. 2 and Supplementary Note), the *attenuation ratio*—(LDSC intercept - 1) / (mean *χ*^2^ – 1)—matched closely between BOLT-LMM and PC-corrected linear regression and was relatively small (Fig. 1c, Supplementary Fig. 2, and Supplementary Table 4). Across 23 traits, we observed similar mean attenuation ratios of 0.078 (s.e.m. 0.006) for PC-corrected linear regression and 0.082 (0.005) for BOLT-LMM, indicating that BOLT-LMM successfully controlled for sample structure (as expected for mixed model methods) [1–3]. (While PC-corrected linear regression on the unrelated British set is not guaranteed to be immune from residual confounding, this sample set has been stringently QC-ed to minimize confounding [5].) In contrast, uncorrected linear regression produced a mean attenuation ratio of 0.104 (0.012), indicating confounding (Supplementary Fig. 2 and Supplementary Table 4). Similarly, PC-corrected linear regression on all European samples exhibited slightly elevated attenuation ratios (mean 0.085, s.e.m. 0.006; binomial *p*=0.01 vs. attenuation on unrelated British samples), indicating slight confounding due to relatedness (Supplementary Table 4), while still achieving lower power than BOLT-LMM (Supplementary Table 2). We note that attenuation ratios are broadly smaller under the LDSC baselineLD model [8], which incorporates functional and LD-related genome annotations, than under the original LDSC model (Supplementary Table 5), consistent with better model fit.

Our new release of the BOLT-LMM software (version 2.3) implements additional computational improvements that reduce running times by a factor of ~4x versus the previous version, achieving run time scaling close to linear in sample size and comparable to linear regression (a few days for UK Biobank analyses; Fig. 1d and Supplementary Table 6). Specifically, BOLT-LMM v2.3 performs much faster processing of imputed genotypes, which we discovered was the bottleneck for analyses of extremely large imputed data sets (e.g., ~93 million variants in the UK Biobank full release). To overcome this bottleneck, we implemented fast multi-threaded support for test statistic computation on imputed genotypes in the new BGEN v1.2 file format (Supplementary Note). Additionally, for analyses of very large data sets, we now recommend including principal component covariates for the purpose of increasing the rate of convergence of the iterative computations performed during BOLT-LMM’s model-fitting steps [3]. Projecting out top PCs improves the conditioning of the matrix computations that BOLT-LMM implicitly performs, roughly halving the number of iterations required for convergence (Supplementary Table 7).

Our results demonstrate the latent power that mixed model association analysis unlocks in very large GWAS, both by reducing the need for sample exclusions and by amplifying effective sample sizes via conditioning on polygenic predictions from genome-wide SNPs. (We note that in general, care must be taken to consider non-additive effects on GWAS and polygenic prediction when retaining related individuals, although we observed here that the level of relatedness in UK Biobank was low enough not to noticeably affect the overall genetic structure of the data set or the interpretation of our results (Supplementary Note).) Our new release of BOLT-LMM makes mixed model association computationally efficient even on extremely large data sets without the need for distributed computing [9]. Our analyses also reveal subtleties in the interpretation of LD score regression intercepts as a means of differentiating polygenicity from confounding in very large GWAS; we suggest that the attenuation ratio may be a more suitable metric as sample sizes continue to increase. Finally, we note two caveats regarding mixed model analysis of binary traits: First, chi-square-based tests (such as BOLT-LMM) can incur inflated type I error rates when used to analyze highly unbalanced case-control traits; here, the binary traits we analyzed were sufficiently balanced for our results not to be impacted by this issue (which begins to arise at case fractions <10% at UK Biobank-scale sample sizes; Supplementary Note and Supplementary Table 8), but in general, the saddlepoint approximation of SAIGE [10] is more robust in such scenarios (although SAIGE currently has the drawbacks of not performing leave-one-chromosome-out (LOCO) analysis or modeling non-infinitesimal genetic architectures, which can reduce its power). Second, conditioning on genome-wide signal can produce loss of power under case-control ascertainment [2,3]; specialized LMM methods are needed for modeling this scenario at scale. Overall, we hope that our findings provide clarity to the GWAS community on analytical best practices for maximizing the value of the UK Biobank and other large biobanks.

## Code and data availability

BOLT-LMM v2.3 is open-source software freely available at http://data.broadinstitute.org/alkesgroup/BOLT-LMM/. Access to the UK Biobank Resource is available via application (http://www.ukbiobank.ac.uk/). BOLT-LMM association statistics computed in this study are currently available for public download at http://data.broadinstitute.org/alkesgroup/UKBB/ and have been submitted to the UK Biobank Data Showcase.

## Acknowledgments

We are grateful to H. Finucane and Y. Reshef for helpful discussions. This research was conducted using the UK Biobank Resource under Application #10438 and was supported by US National Institutes of Health grants R01 HG006399, R01 GM105857, and R01 MH107649 (ALP), a Burroughs Wellcome Fund Career Award at the Scientific Interfaces (PL), and a Boehringer Ingelheim Fonds fellowship (APS). Computational analyses were performed on the Orchestra High Performance Compute Cluster at Harvard Medical School, which is partially supported by grant NCRR 1S10RR028832-01.

